# siRNA containing a unique 5-nucleotide motif acts as a quencher of IFI16-mediated innate immune response

**DOI:** 10.1101/564872

**Authors:** Hongyan Sui, Jun Yang, Xiaojun Hu, Qian Chen, Tomozumi Imamichi

## Abstract

We previously reported that small interfering RNA (siRNA) enhances DNA or DNA virus mediated-interferon (IFN)-λ1 induction through retinoic acid-inducible gene I (RIG-I) and interferon gamma-inducible protein 16 (IFI16) crosstalk-signalling pathway. Here we provide further evidence of a new role for siRNA. siRNA containing a 5-nucleotide (nt) motif sequence suppresses DNA-mediated IFNs and inflammatory cytokines. We determined that motif siRNA inhibits the induction when the motif is located at the 3’ or 5’-terminus of siRNA. Using THP1-Lucia ISG cells with various DNA stimulants, it is demonstrated that motif siRNA inhibits DNA or DNA virus but not RNA virus-mediated signaling. Motif siRNA specifically interrupts IFI16 binding to DNA and has 2.5-fold higher affinity to IFI16 than that of siRNA without the motif. Collectively, these findings may shed lights on a novel function of siRNA with the 5-nt motif as a quencher of innate immunity and facilitate the development of potential therapeutics to treat diseases in which this pathway is dysregulated.

## Introduction

Innate immunity is the first line of defense and relies on germ line–encoded pattern recognition receptors (PRRs) [1]. It has been known for over a decade that DNA, the most recognizable unit of life, is a potent trigger of inflammatory responses in cells [2]. A significant effort from many laboratories has highlighted the importance of cytosolic DNA sensing in the innate immune response against invading DNA focusing on the induction of type-I IFN, Interleukin (IL)-1β or IL-18 [3]. For example, DEAD-box helicase 41 (DDX41) [4], Gamma-interferon-inducible protein (IFI16) [5] and DNA-dependent activator of IFN-regulatory factors (DAI) [6] detect double stranded DNAs (dsDNAs) and activate the STING-TBK1-IRF3 (stimulator of interferon genes - tank-binding kinase 1 - interferon regulatory factor 3) pathway. Leucine-rich repeat flightless-interacting protein 1 (LRRFIP1) binds dsDNA and triggers IRF3 activation through β-catenin [7]. DEAH box protein (DHX)9 and DHX36 associate with dsDNA and lead to NFκB activation through Myeloid Differentiation Primary Response 88 (MyD88) [8, 9]. Cyclic GMP-AMP Synthase (cGAS) has been most recently defined a well-known DNA sensor that recognizes cytoplasmic DNA [10, 11]. Once detect DNA, cGAS undergoes a conformational change that allows ATP and GTP to come into the catalytic pocket, leading to the synthesis of cGAMP, a strong activator of the STING-TBK1 axis [10, 11]. Absent in melanoma 2 (AIM2), a PYHIN (Pyrin-and HIN200-domain-containing) protein, interacts with cytosolic DNA and activates inflammasomes by recruiting apoptosis-associated speck-Like protein (ASC) and pro-caspase-1 and subsequently mature forms of IL-1β and IL-18 were produced [12-14]. Immune responses to DNA are not restricted to type I IFN-inducing pathways: cytosolic DNA also activates type III IFNs [15-19], IFN-λ1 [20-22], IFN-λ2/3 [23, 24] and IFN-λ4 [25, 26]. In human cells, Ku70, a DNA repair protein, predominantly induces IFN-λ1 upon DNA transfection or DNA virus infection and this induction is associated with a translocation of Ku70 from the nucleus to the cytoplasm to bind dsDNA, interacts with STING and promotes production of type III IFNs through activation of IRF3, IRF1 and IRF7 [20, 27]. Of note, the mouse *ifnl1* gene lacks exon 2 and contains a stop codon within exon 1, which results in the failure to encode a functional IFN-λ1 protein [16]. So, the study related to a physiological relevance of the induction of IFN-λ1 in a mouse model is not capable.

Investigation of the mechanism of the Innate immune activation by cytosolic DNA from microbial pathogens has been extensively studied in terms of both DNA binding proteins and the mechanism of DNA recognition for subsequent downstream signaling and immune activation [28, 29]. However, in some scenarios, those cascade inductions of cytokines or inflammatory chemokines maybe harmful for the host cells, some diseases are highly related with those over-responses of innate immunities [30]. Some reagents must possess mechanisms to subvert the signaling pathway to establish successful regulation. However, this kind of study is rarely reported.

Small interfering RNAs (siRNAs) are artificially synthesized 19–23 nucleotide long double-stranded RNA molecules [31]. They are routinely used in molecular biology for a transient silencing of gene of interest. Except the biological function of specific gene knocking down by siRNA, other potential functions of siRNA are also reported. siRNAs were originally thought incapable of inducing immune responses because they are short and designed to mimic the natural product of Dicer *in vivo*. However, many studies have reported innate immune stimulation by siRNA and/or the siRNA delivery vehicle [32-35]. Various features of the siRNA structure, sequence, and delivery mode have contributed to the immune stimulation effect, leading to undesired effects and the misinterpretation of experimental results, for example the production of IFN-α, tumor necrosis factor alpha (TNF-α), IL-6 and IL-12 [32, 36-39]. Previous studies indicated that transfection of siRNA could activate double-stranded RNA-activated protein kinase R (PKR) [40] or TLR3 [41] in a sequence-independent manner or sequence-dependently recognized by TLR7 in mice and TLR7/8 in humans [42]. And our recent study also consistently indicated that siRNA cooperated with invading DNA by DNA transfection or DNA virus infection, highly induces type III IFN response through RIG-1 and IFI16 crosstalk signaling pathway. This finding further shed light on the widely existed but uncovered functions of siRNA [21]. Coincidence with our previous study, we reported here, another new function of siRNAs. Except for the RNA interference and innate immunostimulatory function of siRNA, we found siRNA containing a unique 5-nt motif at the 3’ or 5’ terminus of siRNA sense strand potently inhibits a DNA sensor, IFI16-mediated innate immune response in response to DNA or DNA virus-derived intracellular stimulation. The finding may provide a new strategy to regulate the over-response of the innate immune system for the host cells upon pathogen invasion, shedding light on a novel function of siRNA with the unique 5-nt motif as a quencher of the innate immunity and a potential therapeutic agent for auto immune diseases.

## Results

### The impact of different siRNA transfection on DNA-mediated *IFNL1* induction

We previously reported that siRNA enhances DNA transfection or DNA virus infection-mediated type III IFN response through a crosstalk between RIG-I (an RNA sensor) and IFI16 (a DNA sensor) signaling pathway [21]. In a course of study, to identify a mechanism of the siRNA-mediated *IFNL1* induction in IFN-α-treated Hela cells, a total of 48 siRNAs including non-target siRNAs (Ctrl-siRNA, siRNA targeting GFP or luciferase) and siRNAs targeting IFN-stimulatory genes (ISGs) were transfected into the IFN-α-treated HeLa cells, and the cells were further stimulated by DNA transfection to induce DNA-mediated innate immune response. The cells were then collected 6 h after DNA transfection to detect *IFNL1* mRNA expression level by real-time RT-PCR. The experimental procedure was presented in the Figure 1A. The results consistently showed that some siRNAs highly enhanced DNA-mediated *IFNL1* induction, for example Ctrl-siRNA, siIFIT1-1, siTrim25, siIFIT5-2, siDENND2D-1, siDENND2D-2 and siPCGF1-1 (Figure 1B & 1C). The sequences of all the siRNAs to enhance DNA-mediated signaling were listed in the Table EV1. Simultaneously, we were also noticed that some of two different siRNAs targeting same gene but encoding different seeding RNA sequence demonstrated a conflicting result, one siRNA enhanced or had no effect, and one inhibited the DNA-induced *IFNL1* induction. They were siRNAs targeting IFIT1 (siIFIT1-1 and siIFIT1-2), siRNAs targeting TRIM22 (siTRIM22-1 and siTRIM22-2) and siRNAs targeting LOC732371 (siLOC732371-1 and siLOC732371-2). Their sequences of those three siRNAs which inhibited DNA-induced *IFNL1* induction was listed in the Figure 1D. A Sequence analysis illustrated that one 5-nucleotide (nt) sequence 5’-AAGAA-3’ was found in all three sense strands of siRNAs, we call it as 5-nt motif in our study.

**Figure 1.**
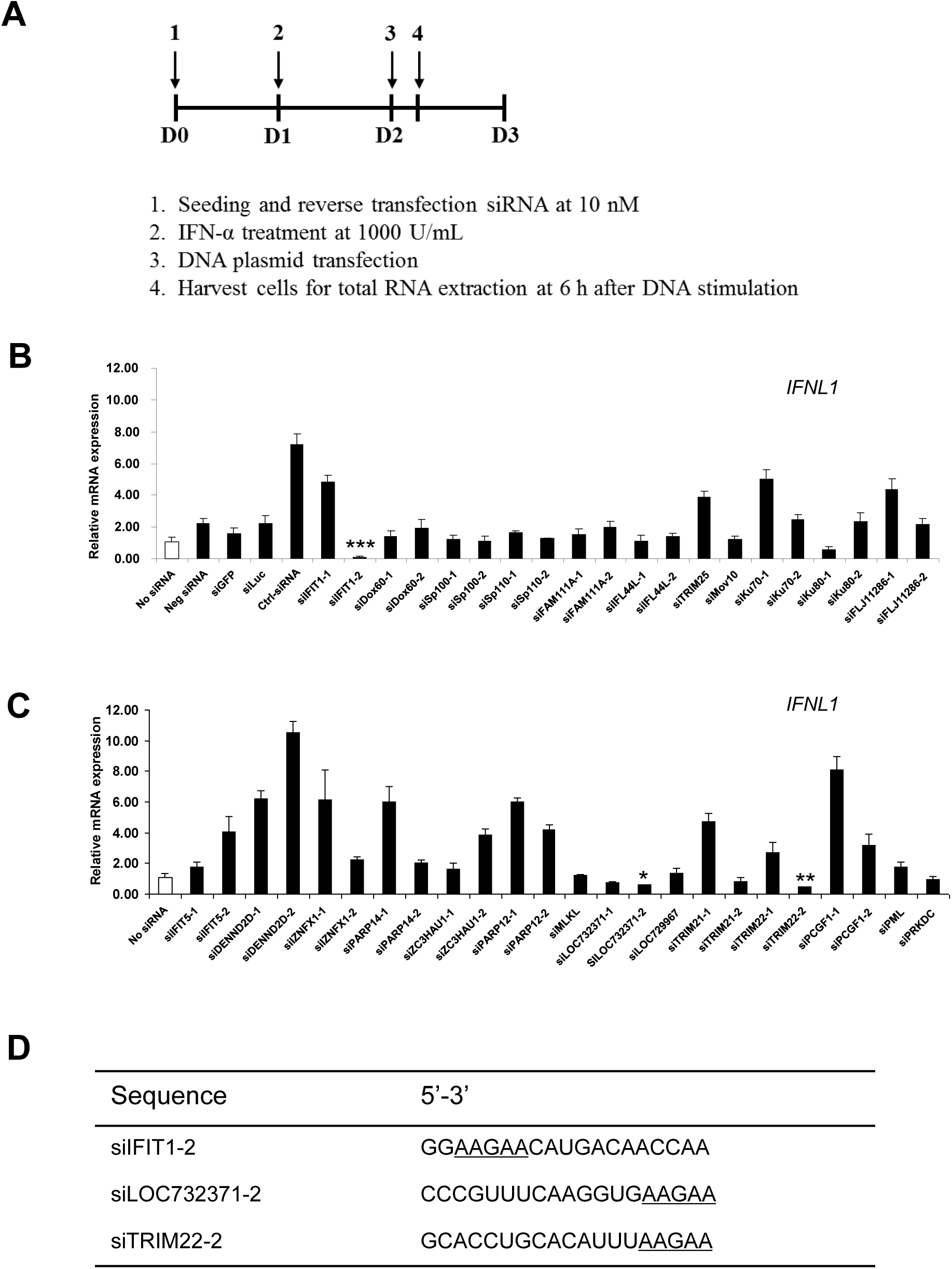
The impact of siRNA transfection on DNA-mediated *IFNL1* production. (**A**) an experimental flowchart. Briefly, HeLa cells were transfected with 10 nM siRNA, and then treated with IFN-α 24 h after siRNA transfection. On day 3, the cells were further transfected by DNA. Total RNA was extracted 6 h after DNA transfection. (**B**, **C**) Relative *IFNL1* expression was measured by real time RT-PCR, and the gene expression level was compared with the cells with DNA transfection but no siRNA transfection. All results represent the mean of three independent experiments. Error bars show SD. \**p* < 0.05, \*\**p* < 0.001, \*\*\**p* < 0.0001 (Student’s *t* test) was indicated in the figure where the gene expression level was compared with the cells with DNA transfection but no siRNA transfection. (**D**) the motif 5’-AAGAA-3’ exits in all three siRNAs which inhibit DNA-mediated *IFNL1* induction.

### Motif siRNA motif-location dependently affects DNA-mediated *IFNL1* induction

To assess whether this 5-nt motif sequence was responsible for suppressing the DNA-mediated *IFNL1* induction, we used Ctrl-siRNA which highly enhances DNA-mediated IFN response as a control sequence [21]. Total 6 variants of siRNAs that encoded the motif sequence at different location in the RNA sequence were designed and chemically synthesized, as shown in the Figure 2A. The original sequence of Ctrl-siRNA (wild type) was replaced with 5-nt motif sequence; the motif was located from the 5’ terminus to 3’ terminus of siRNA sense strand and was represented by 5’m-siRNA, 5’m1-siRNA, 5’m2-siRNA, mm-siRNA, 3’m1-siRNA and 3’m-siRNA respectively (Figure 2A). And then following the same experimental method as shown in Figure 1A, those variants were transfected into HeLa cells, and followed by IFN-α treatment on day 2 and DNA transfection on day 3. The cells were subjected by total cellular RNA extraction at 6 h or 24 h after DNA transfection, and the levels of *IFNL1* mRNA expression were measured by real time RT-PCR (Figure 2B). The results showed that the impact of 6 motif siRNAs on *IFNL1* induction was different. Variants with the motif either at 3’ terminus or 5’ terminus (5’m-siRNA and 3’m-siRNA) showed no effect on the IFN induction compared with the cells without siRNA transfection. And the variant with the motif in the middle (mm-siRNA) indicated a strong enhancement for the IFN induction as does Ctrl-siRNA. Collectively, the impact of motif siRNA on the DNA-mediated IFN induction was shown a bell-shaped response along with changes of the motif location from 5’ terminus to the 3’ terminus. and the results from the samples collected at 24 h after DNA transfection were consistent with that of 6 h time point. In summary, the motif siRNA affected DNA-mediated *IFNL1* induction with a location dependent manner.

**Figure 2.**
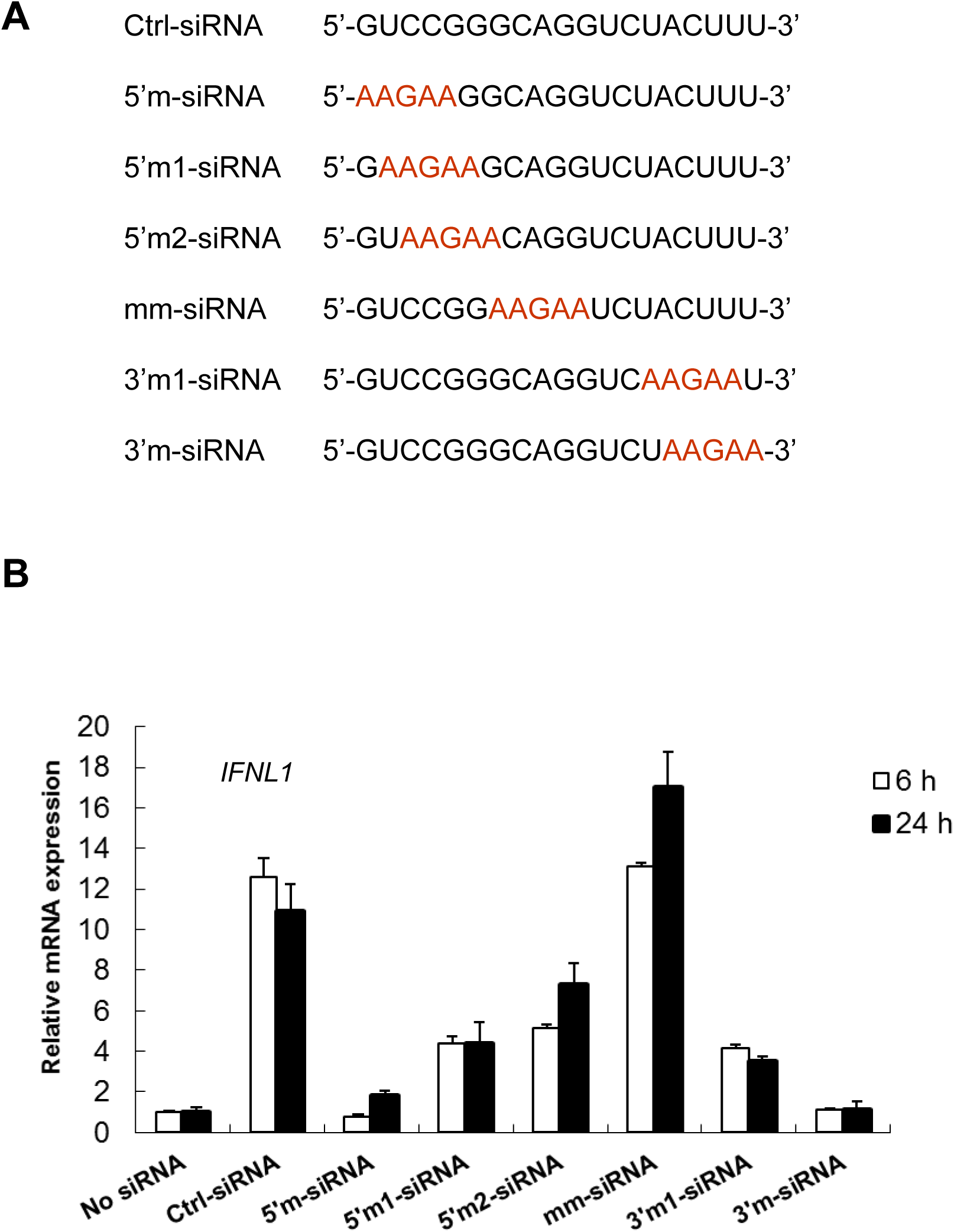
Motif siRNA motif-location dependently affects DNA-mediated *IFNL1* induction. (**A**) the sense strand sequence of Ctrl-siRNA which highly enhances DNA-mediated *IFNL1* induction. The design of other siRNAs with motif sequence, which locates from the 5’ terminus to the 3’ terminus of the sense strand. (**B**) HeLa cells were transfected with 10 nM motif siRNA and then treated with IFN-α 24 h after siRNA transfection. On day 3, the cells were further transfected by plasmid DNA. Total RNA was extracted 6 h or 24 h after DNA transfection, relative *IFNL1* expression was measured by real-time RT-PCR, and the gene expression level was compared with the cells with DNA transfection but no siRNA transfection.

### Motif siRNA potently suppresses Ctrl-siRNA-enhanced DNA-mediated IFNs and inflammatory cytokines

To further clarify a role of the inhibitory effect of motif siRNA, different dose of 3’m-siRNA or mm-siRNA was co-transfected with 10 nM Ctrl-siRNA in HeLa cells, and then followed by IFN-α treatment on day 2 and DNA transfection on day 3. The *IFNL1* mRNA expression level was measured by real time RT-PCR (Figure 3A). The results showed that 3’m-siRNA dose-dependently inhibited the Ctrl-siRNA/DNA-mediated *IFNL1* induction, in contrast, the variant encoding the motif sequence in the middle (mm-siRNA) did not show the similar suppression. This result indicated that the 3’m-siRNA was able to compete Ctrl-siRNA-enhanced signal in the IFN-induction. To further define whether the 3’m-siRNA suppresses other cytokine induction, gene activation of *IFNL2/3, IFNB, CXCL10, RANTES* and *TNFA* were also analyzed. The results clearly indicated that co-transfection of 3’m-siRNA suppressed the Ctrl-siRNA/DNA-mediated cytokine inductions (Figure 3B). Therefore, motif siRNA with motif at an end location was able to inhibit the production of not only IFN but also inflammatory cytokines.

**Figure 3.**
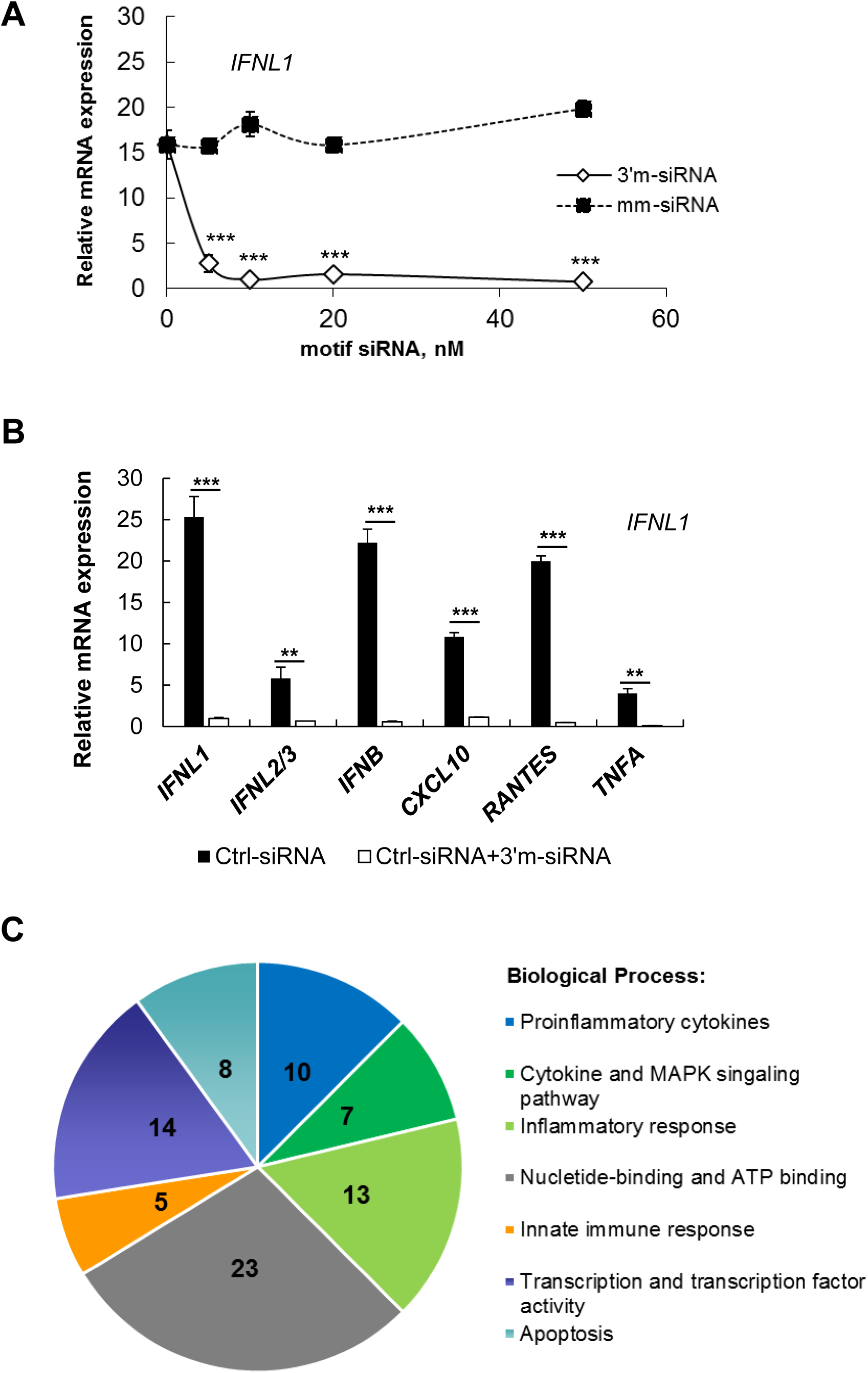
Motif siRNA 3’m-siRNA dose-dependently inhibits Ctrl-siRNA-enhanced, DNA-mediated IFNs and inflammatory cytokines. (**A**) HeLa cells were transfected with 10 nM Ctrl-siRNA and different dose of motif siRNAs, mm’-siRNA or 3’m-siRNA. And then treated with IFN-α 24 h after siRNA transfection. On day 3, the cells were further transfected by plasmid DNA. Total RNA was extracted 6 h after DNA transfection, relative *IFNL1* expression was measured by real-time RT-PCR, and the gene expression level was compared with the cells with DNA transfection but no siRNA transfection. \*\*\**p* < 0.0001 (Student’s *t* test) was indicated in the figure to show the significance difference between 3’m-siRNA and mm-siRNA at a same dose treatment. (**B**) HeLa cells were transfected with 10 nM Ctrl-siRNA or a combination of 10nM 3’m-siRNA and 10nM Ctrl-siRNA, and then treated with IFN-α 24 h after siRNA transfection. On day 3, the cells were further transfected by plasmid DNA. Total RNA was extracted 6 h after DNA transfection, relative *IFNL1, IFNL2/3, IFNB, CXCL10, RANTES* and *TNFA* expression was measured by real-time RT-PCR, and the gene expression level was compared with the cells with DNA transfection but no siRNA transfection. \*\**p* < 0.001, \*\*\**p* < 0.0001 (Student’s *t* test) was indicated in the figure. (**C**) Genes involved in immunity and inflammatory cytokines were downregulated by 3’m-siRNA. The 113 genes were enriched ≥1.5-fold in cells transfected with Ctrl-siRNA relative to a combination of Ctrl-siRNA and 3’m-siRNA on the microarray. Those genes were subject to gene ontology analyses (DAVID) and classified according to biological process (P ≤ 0.05). The set of 51 genes among 113 that belong to each class was given.

To further characterize a gene regulation profile in the cells transfected with the 3’m-siRNA, a microarray analysis was performed. Following the same experimental protocol, we transfected HeLa cells with 10 nM Ctrl-siRNA alone or 10 nM Ctrl-siRNA and 3’m-siRNA, and then harvested RNA at 6 h after DNA transfection. The gene expression was compared between presence and absence of 3’m-siRNA. From three independent experiments, we found that the expression of 113 genes was ≥1.5-fold greater with Ctrl-siRNA than with the combination of 3’m-siRNA and Ctrl-siRNA. Using gene ontology analyses (DAVID), we found that the set of 51 genes was significantly enriched for genes involved in biological processes such as innate immune response and inflammatory cytokines (Figure 3C). The details for those 51 genes were provided in the Table EV2. This observation was consistent with the idea that motif siRNA with the motif at the 3’ or 5’ terminus suppressed the induction of genes involved in the interferon and inflammatory response. Several genes involved in apoptosis were also presented. These data confirm that motif siRNA with the motif at the 3’ or 5’ terminus specifically suppressed the induction of genes involved in innate immune responses and some cytokine induction pathways.

### Exchange of any nucleotide of the motif sequence decreases the inhibitory efficacy of 3’m-AAGAA siRNA

Through screening the impact of 48 siRNAs on IFN induction, we found that the motif 5’-AAGAA-3’ existed in all siRNAs which inhibited DNA transfection-induced *IFNL1* induction. And this motif must be located at the end terminus of the siRNA to make motif siRNA potently inhibited Ctrl-siRNA/DNA-induced interferon response and inflammatory cytokines. Those observations led us to speculate whether the 5-nt sequence is essential for motif siRNA to inhibit DNA-induced signaling. So, we further designed several motif siRNAs with one nucleotide change and compared the inhibitory effect among the variants. The designed mutants are listed in the Figure 4A. And then those motif siRNAs were transfected to HeLa cells with 10 nM Ctrl-siRNA to define whether the motif siRNA still inhibited Ctrl-siRNA/DNA-mediated *IFNL1* induction. Compared with 3’m-AAGAA siRNA, all other motif siRNA with one nucleotide change, 3’m-AA<u>A</u>AA, 3’m-AA<u>U</u>AA, 3’m-AA<u>C</u>AA, 3’m-A<u>C</u>GAA, 3’m-AAG<u>U</u>A and 3’m-AAGA<u>U</u> increased *IFNL1* induction. The results indicated that the inhibitory activity of 5-nt AAGAA motif siRNA is depends on the sequence of the motif. We further confirmed here that the immunosuppressive motif sequence is 5’-AAGAA-3’.

**Figure 4.**
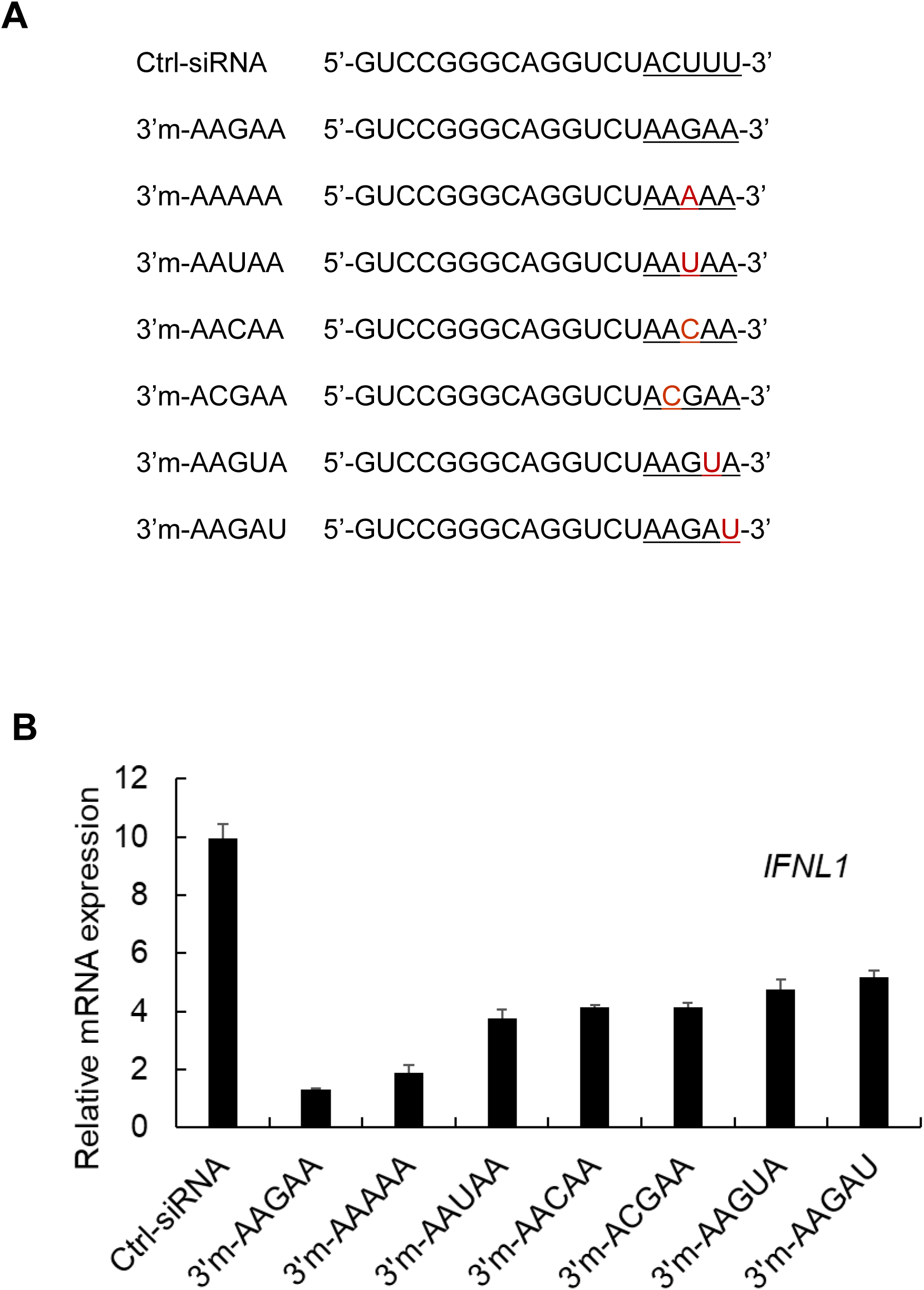
Exchange any nucleotide of the motif sequence decreases the inhibitory efficacy of 3’m-AAGAA motif siRNA. (**A**) the list of 3’m-siRNA sense strand (5’-3’) with one nucleotide change compared with original 3’m-AAGAA sequence. (**B**) HeLa cells were transfected with 20 nM indicated motif siRNA with or without 10 nM Ctrl-siRNA and then treated with IFN-α 24 h after siRNA transfection. On day 3, the cells were further transfected by plasmid DNA. Total RNA was extracted 6 h after DNA transfection, relative *IFNL1* expression was measured by real-time RT-PCR, and the gene expression level was compared with the cells with DNA transfection but no siRNA transfection.

### Motif siRNA suppresses DNA or DNA virus-mediated innate immune response

As the motif siRNA abolished DNA plasmid-mediated IFN induction, we also examined whether the motif siRNA inhibits other stimulants-induced innate immune response. This experiment was performed using a THP1-Lucia ISG cells. THP1-Lucia ISG cells were derived from the human monocytic cell line THP-1, which represents a model of choice to study the activation and signaling of Cytosolic DNA Sensors (CDS), and THP-1 cells have been shown to express all the CDS identified so far [4, 43, 44], with the exception of DAI [6, 45]. Using this cells as a screening platform, we transfected 3’m-siRNA at a dose of 0, 10, 20 and 50 nM, and then the cells were stimulated by different stimulants for CDS, Plasmid DNA, poly(A:T), 2’3’-cGAMP, VACV-70 and HSV-60, Lucia luciferase in the cells was monitored as an indicator for immune response. the Ctrl-siRNA-enhanced DNA-mediated immune response was also included in this experiment as a positive control. The result demonstrated that all the tested stimulants initiated a certain level of luciferase activity, and the induction of Luciferase activity by all CDS agonists but 2’3’-cGAMP was decreased by a motif siRNA-dose dependently manner (Figure 5A). Furthermore, we tested whether motif siRNA suppresses virus-mediated innate immune response in THP1 cells. the THP1 cells were first transfected by 3’m-siRNA at different doses. and then the cells were infected by a DNA virus, HSV-1 at a MOI of 5 (Figure 5B) or an RNA virus, SeV at a final concentration of 50 HA units/mL (Figure 5C). The RNA was extracted from the cells at 24 h after virus infection. The *IFNL1* mRNA expression level was detected by real-time RT-PCR. The results showed that 3’m-siRNA dose dependently suppressed HSV-1 virus, but not SeV virus-mediated *IFNL1* induction (Figure 5B and 5C). Taken together, the motif siRNA inhibited only DNA (DNA ligands or DNA virus)-mediated innate immune response.

**Figure 5.**
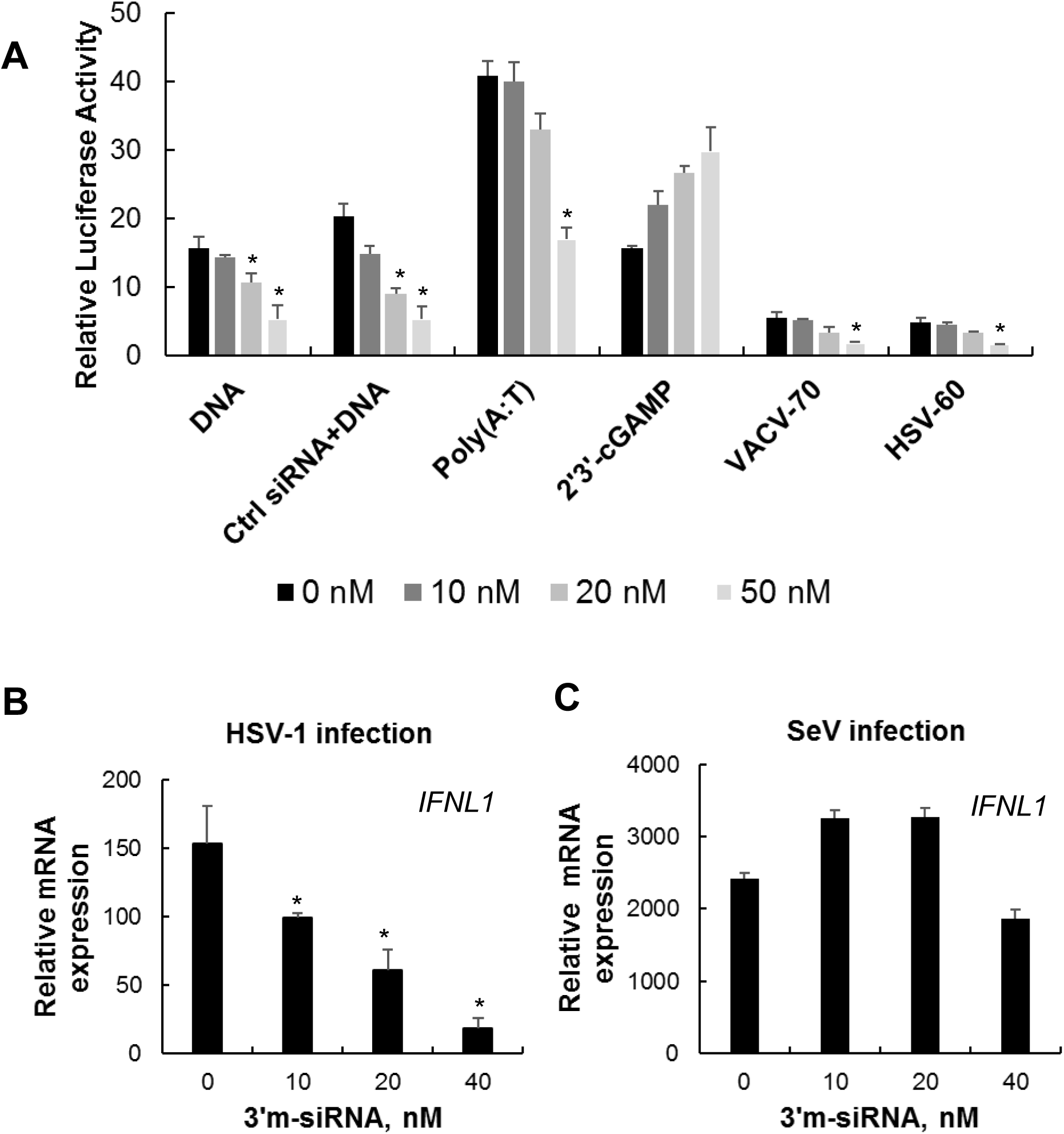
Motif siRNA inhibits DNA or DNA virus–induced innate immune response. (**A**) THP1-Lucia ISG cells were transfected with different dose of 3’m-siRNA and then further treated with various DNA stimulants. The immune response indicated by Lucia luciferase was detected 24 h after stimulation. \**p* < 0.05 (Student’s *t* test) was indicated in the figure to show the significance difference when compared with the same DNA stimulant alone-induced luciferase activity. (**B, C**) THP1 cells were transfected with different dose of 3’m-siRNA and then infected by (**B**) HSV-1 virus at a MOI of 5 or (**C**) Sendai virus at 40 HA unites/mL. Total RNA was extracted 24 h after virus infection, relative *IFNL1* expression was measured by real-time RT-PCR, and the gene expression level was compared with untreated cells. \**p* < 0.05 (Student’s *t* test) was indicated in the figure where the gene expression level was compared with the cells with HSV-1 virus infection-induced *IFNL1* induction.

### Motif siRNA dose-dependently inhibits HSV-1 virus infection-mediated innate immune response in human primary immature dendritic cells

To confirm a physiological relevance of the finding, we further assess the impact of motif siRNA transfection in DNA virus-mediated innate immune response in primary immature dendritic cells (iDCs). iDCs cells were transfected with different amounts of 3’m-siRNA and then cells were infected with HSV-1 at a MOI of 5. The cells were collected to measure *IFNL1, IFNL2/3, IFNB, CXCL10, RANTES* and *TNFA* mRNA expression. The results consistently showed that motif siRNA does-dependently inhibited HSV-1-mediated not only *IFNL1, IFNL2/3, IFNB*, but also inflammatory cytokines, *CXCL10, RANTES* and *IFNA* (Figure 6A, 6B, 6C, 6D, 6E and 6F respectively). Therefore, the results from human primary cells further emphasized that an insight of motif siRNA as a potent therapeutic reagent to regulate auto immune diseases.

**Figure 6.**
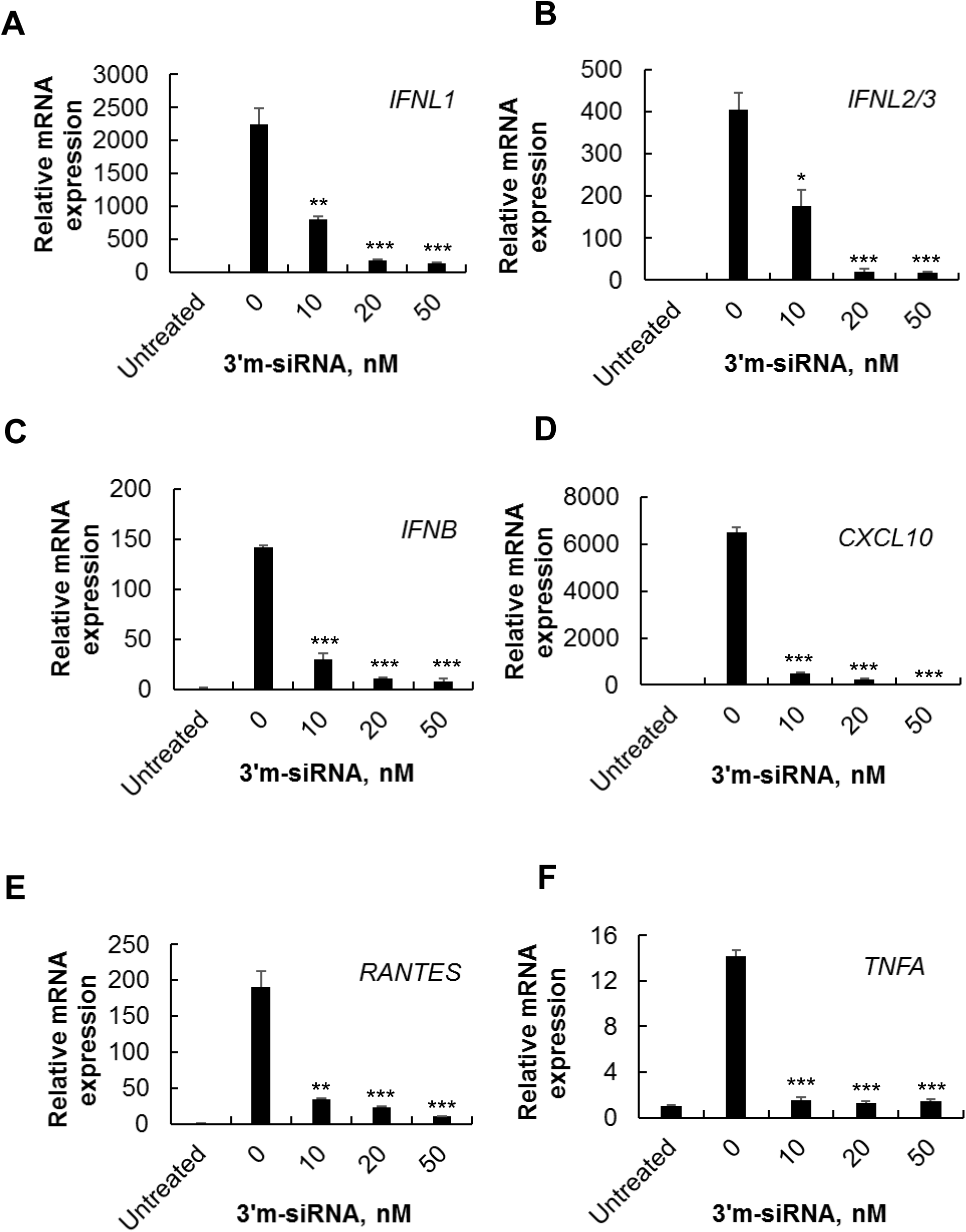
Motif siRNA dose-dependently inhibits HSV-1 virus infection-mediated innate immune response in human primary immature dendric cells. Immature dendritic cells (iDC) were produced from monocytes isolated from healthy donors. iDC were first transfected by 3’m-siRNA at indicated concentrations, and one day later were infected by HSV-1 at a MOI of 5, the cells were collected for RNA extraction 24 h after virus infection. Relative (**A**) *IFNL1*, (**B**) *IFNL2/3*, (**C**) *IFNB*, (**D**) *CXCL10*, (**E**) *RANTES* and (**F**) *TNFA* mRNA expression was measured by real-time RT-PCR, and the gene expression level was compared with untreated cells. \**p* < 0.05, \*\**p* < 0.001, \*\*\**p* < 0.0001 (Student’s *t* test) was indicated in the figure where the gene expression level was compared with the cells with HSV-1 infection alone-induced gene expression levels.

### Motif siRNA specifically interrupts the binding of DNA to IFI16

To dissect the inhibitory effect of siRNA containing the motif at 3’ or 5’ terminus, we examined the binding affinity of the motif siRNA to a DNA sensor protein using a DNA-conjugated pull-down assay with or without a competition strategy, the experimental protocol was shown in the Figure 7A. First, we performed a DNA-conjugated pull-down assay to define if motif siRNA interrupted DNA binding to specific DNA sensor. Biotinylated-DNA conjugated streptavidin beads were incubated with cytoplasm fraction in the absence or presence of various siRNA competitors. After incubation, proteins bounded to the beads were separated on SDS-PAGE and detected by western blot. In the pull-down assay, we added 10X higher amount of Ctrl-siRNA, mm-siRNA or 3’m-siRNA as a competitor. The results indicated that DNA binding to DNA sensor IFI16 was not interrupted when Ctrl-siRNA or mm-siRNA was added as a competitor. However, we found that the band of IFI16 was diminished in the presence of the competitor, 3’m-siRNA (Figure 7B). Therefore 3’m-siRNA interrupts the binding of DNA to IFI16. To elucidate whether the interruption by 3’m-siRNA is unique to IFI16, the same pull-down assay was performed using cytosol fraction from THP-1 cells which contains multiple DNA sensors including cGAS and Ku 70 and the bounded proteins were detected by anti-cGAS or Ku70 antibody. As shown in the Figure 7C, the data indicated that 3’m-siRNA as a competitor had no interruption with DNA binding to cGAS or Ku70. Therefore, motif siRNA with the motif at the end terminus interrupts the binding of DNA to IFI16 and has no impact on the DNA sensing of cGAS or Ku70.

**Figure 7.**
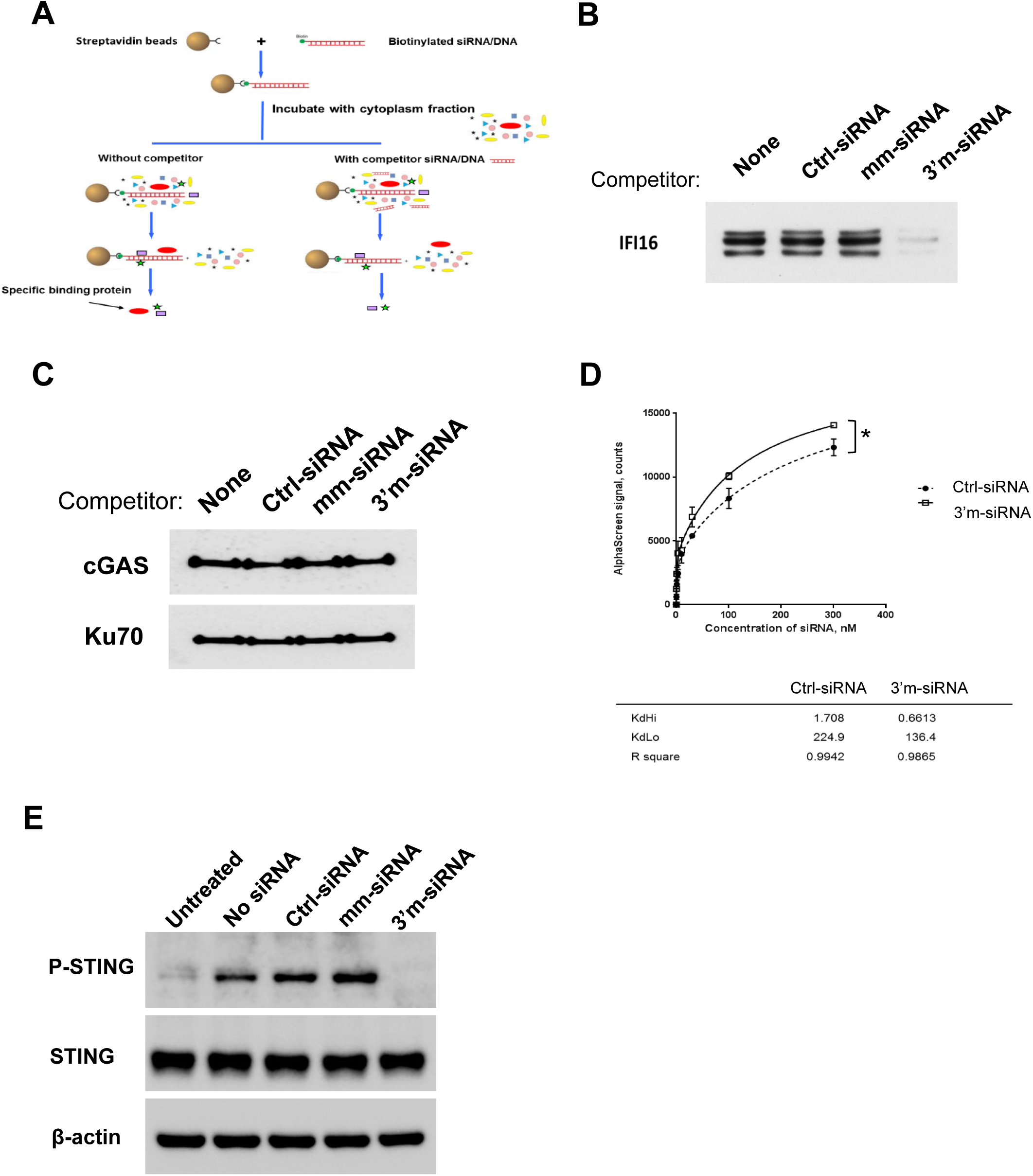
Motif siRNA interrupts the binding of DNA to IFI16, therefore inhibiting activation of the downstream signaling. (**A**) a diagram view of nucleotide pull-down assay with or without a competitor. (**B**) A set of DNA pull-down assay was performed in the absence or presence of Ctrl-siRNA, mm-siRNA or 3’m-siRNA as a competitor. Bound proteins on the beads were separated by SDS-PAGE, and Western blot was performed using anti-IFI16 antibody. (**C**) 3’m-siRNA has no interruption with DNA binding to other DNA sensor protein, cGAS or Ku70. A set of DNA pull-down assay was performed in the absence or presence of Ctrl-siRNA, mm-siRNA or 3’m-siRNA as a competitor. Bound proteins on the beads were separated by SDS-PAGE, and western blot was performed using anti-cGAS and anti-Ku70 antibodies. (**D**) An AlphaScreen assay was performed to estimate the binding affinity between IFI16 and siRNA molecules. IFI16-overexpressed 293T cell lysate was incubated with different concentrations of siRNA molecules bearing a biotin at 5’ end of the antisense strand. The dissociation constant Kd is calculated by statistical analysis (nonlinear regression). The *p* values between two groups is 0.0038, and \**p* < 0.05 (Student’s *t* test) was indicated in the figure. (**E**) 3’m-siRNA inhibited IFI16-mediated downstream signaling, the phosphorylation of STING. THP-1 cells were transfected with different siRNAs, Ctrl-siRNA, mm-siRNA and 3’m-siRNA. And the cells were further treated by DNA transfection. The cell lysate was collected 1h after DNA transfection and Western blot was performed using anti-p-STING, STING and β-actin antibodies.

To compare the binding affinity between Ctrl-siRNA and 3’m-siRNA to IFI16, we performed an AlphaScreen assay. The cell lysates from the His-tagged IFI16 overexpressed 293T cells were incubated with different concentrations of 5’-biotinylated Ctrl-siRNA or 3’m-siRNA. The resulting florescence signal correlates with the number and proximity of interacting donor (Biotin-siRNA)-acceptor (His-IFI16) pairs. The concentration of indicated siRNAs is plotted against the Alpha Screen signal (Figure 7D). Dissociation constant (kd) was calculated using the Prism. There was an inverse relationship between Kd and binding affinity. The calculated results showed that both KdHi and KdLo value of Ctrl-siRNA were higher than those of 3’m-siRNA. KdHi and KdLo for Ctrl-siRNA is 1.708 and 224.9, While KdHi and KdLo for motif siRNA is 0.6613 and 136.4, respectively, suggesting that the binding affinity of motif siRNA to IFI16 is 2.5-fold greater than Ctrl-siRNA (p=0.038). The AlphaScreen assay further supported the results in the figure 7B. It was reasonable to consider that siRNA with motif at 3’ or 5’ terminus had a higher binding affinity to IFI16, therefore blocked IFI16 binding to invaded DNA.

It is reported that IFI16 senses DNA and then induce phosphorylation of STING [5]. Therefore, we further assessed whether the downstream signaling of IFI16 is inhibited by 3’m-siRNA. We transfected THP-1 cells with different siRNAs, Ctrl-siRNA, 3’m-siRNA or mm-siRNA. And then the cells were further stimulated by DNA transfection. The whole cell lysate was collected at 1h after DNA transfection. Western blot was performed to detect the phosphorylation of STING, an important activation marker at downstream of IFI16. The result, as shown in the figure 7E, suggested that the phosphorylation of STING was inhibited by 3’m-siRNA, but not by mm-siRNA or Ctrl-siRNA. This result consisted with that 3’m-siRNA interrupts the binding of DNA to IFI16, and therefore inhibiting the downstream signaling for IFN response.

## Discussion

siRNA was thought too short to stimulate innate immunity. However, several groups found that siRNAs can induce type I IFN through various RNA sensors, RIG-1, PKR, TLR3 and TLR7/8 in human primary cells and many human cell lines [33, 46-50]. We have previously reported that siRNA transfection enhances DNA-induced IFN-λ1 (type-III IFN) induction in human cells and found that the augmentation was caused by a cross talk between RIG-I and IFI16 [21]. In the current study, we provided a novel function of siRNAs: siRNA containing a 5-nt motif sequence AAGAA at either 3’ or 5’ terminus (3’m or 5’m-siRNA) acts as a quencher for IFI16-mediated IFNs and inflammatory cytokines.

Among screened 48 siRNA in this study, we found three siRNAs suppressed DNA-induced innate immune response. The results indicated that the inhibition was not mediated by the product of the siRNA-targeting gene, instead, the inhibition may be caused in a siRNA sequence dependent manner. We compared sequence similarity among those three siRNAs, and then we found a 5-nt sequence commonly existed in all three siRNAs. This AAGAA sequence was essential for inhibitory activity of motif siRNA, since with any one nucleotide change decreased the immunosuppressive efficacy of 3’m-AAGAA siRNA. Several groups have found some motifs in the context of siRNA sequence which have impact on the immunostimulatory characteristics of siRNAs. Immunorecognition of RNA depends on certain molecular features such as length, double-versus single-stranded configuration, sequence motifs, and nucleoside modifications such as triphosphate residues. Judge and colleagues compared cytokine induction by transfection of 16 different synthetic siRNAs against GAPDH or a gene called BP1, a breast cancer–associated protein in PBMC, and identified the motif 5′-UGUGU-3′ responsible for cytokine induction [51, 52]. Similar with those work, Hornung et al. found that siRNA can induce robust IFN-α in pDC with siRNA sequence dependent manner. They defined the motif 5′-GUCCUUCAA-3′ responsible for IFN-α induction [42]. Several years ago, Zheng and his colleagues demonstrated that a group of highly effective siRNAs targeting different influenza virus H5N1 strain shares a unique motif, 5’-GGAGU-3’ and demonstrated that the structure of the unique motif is critical in determining the potency of the siRNA-mediated inhibition against viral infection. And using a mouse model, they demonstrated that the viral suppression is associated with early productions of β-defensin and IL-6 production by the motif siRNA [53]. In our current study, we found that the motif 5’-AAGAA-3’ negatively regulated type III IFN induction in human cells. The immunosuppressive motif sequence is distinct from all the reported motifs so far.

Further characterization clarified that the suppression effect of motif siRNA depends on the location of the motif. We designed several motif siRNAs located at different location from 5’ terminus to 3’ terminus. the screening suggested that the 5-nt motif must be located at 3’ or 5’ terminus, and then the motif siRNA has the capability to inhibit DNA-mediated signaling. The suppressing effect of the motif is completely diminished when the motif sequence in the middle of siRNA. A precise mechanism related with the location of motif remains unclear. An RNA structure analysis of the motif siRNA, as shown in the Figure EV1, indicated that mm-siRNA forms a round closed structure, this may make mm-siRNA difficult to access the binding site of DNA sensor IFI16. By contrast, 3’m-or 5’m-siRNA possesses a stem-loop structure with the motif sequence close to the open end, suggesting that all those structures may provide a certain flexibility for motif siRNA binding to IFI16. Even Ctrl-siRNA has a similar folding structure with that of 3’m-siRNA, but this siRNA doesn’t contain motif sequence, indicating that motif sequence itself makes a difference in binding affinity of the siRNA to IFI16, it may be caused in a sequence dependent fashion. Further study regarding the modeling of motif siRNA binding to DNA sensor IFI16 may provide more valuable insights.

For the function of the motif siRNA, we further confirmed that motif siRNA inhibited not only DNA-induced type III interferon response but also the production of IFNα, IFNβ and inflammatory cytokines, RANTES, TNFα. Overall, we demonstrated *in vitro* that motif siRNA dose-dependently quenches DNA-induced innate immune cascades. It is validated that motif siRNA down-regulated the expression of numerous host defense genes, including pattern recognition receptors, kinases, transcription regulators, cytokines, chemokines, apoptosis as assessed by genome-wide microarray analyses. As It is known that dendric cells are an important source of cytokines [54]. Transfection of motif siRNA caused a marked reduction of cytokine and chemokine production induced by human primary iDCs after exposure to DNA virus, HSV-1 infection. Those data provide further physiological relevance of our finding. Similar to our study, Thierry Roger et al. reported that histone deacetylase inhibitors impair innate immune responses to toll-like receptor agonists and to infection [55]. They identified an essential role for acetylation of histones and nonhistone proteins in the regulation of inflammatory and innate immune gene expression, and in host defensive responses against microbes. In an earlier study by Patrice Vitail and his colleagues, IU-dsRNA suppresses the induction of interferon-stimulated genes (ISGs) and apoptosis by poly(I:C). This is another form of innate immune response suppressor [56]. And recently Wu Jian-Jun et al. reported that Kaposi’s sarcoma-associated herpesvirus (KSHV) ORF52, an abundant gamma herpesvirus-specific tegument protein, subverts cytosolic DNA sensing by directly inhibiting cGAS enzymatic activity. They revealed a mechanism through which gamma herpesviruses antagonize host cGAS DNA sensing [57]. In another way, our finding in the current study validated a new function of siRNA with a unique 5-nt motif potently inhibits IFI16-mediated innate immune IFNs and proinflammatory cytokines in response to DNA or DNA virus invasion.

Several arguments led us to believe that these observations may have clinical implications. To further clarify what agonist-mediated innate immune response will be potentially inhibited by motif siRNAs, we established a cell culture model using THP1-Lucia ISG cells, which is derived from THP-1 cells, and has most of reported DNA sensor proteins except DAI [58]. We defined that motif siRNA suppressed DNA-plasmid, DNA together with immunostimulatory siRNA, Poly(A:T), VACV-70 and HSV-60 mediated signaling, but not 2’3’-cGAMP-mediated innate immune response. This result led us speculated that motif siRNA may inhibit the step of DNA binding to DNA sensors. 2’3’-cGAMP is produced in mammalian cells by cGAS in response to double-stranded DNA in the cytoplasm, serves as a second messenger to activate innate immune responses by binding to STING and subsequently inducing the TBK1-IRF3-dependent production of IFN-β [11]. So, we believe motif siRNA is not targeting to inhibit the downstream signaling of DNA sensors. We further validated that motif siRNA does-dependently dampened down HSV-1, a DNA virus, mediated innate immune response, but not SeV virus, an RNA virus. Taken together, the motif siRNA only inhibited DNA or DNA-virus mediated innate immune response and has no effect on RNA virus-mediated signaling transduction.

Further DNA pull-down assay and AlphaScreen analysis revealed that motif siRNA competes binding to DNA sensor IFI16, therefore it interrupts the binding of DNA or DNA virus to IFI16. This interruption is specific for IFI16. Using the similar strategy, motif siRNA cannot compete bind with Ku70 or cGAS in THP-1 cells. Taken together, those results suggest that motif siRNA only suppress IFI16-mediated innate immune response to intracellular DNA or DNA virus stimulation. The motif siRNA has a 2.5 folds stronger binding affinity to IFI16 than Ctrl-siRNA. Wu jian-jun and his colleagues reported that ORF52, a protein in KSHV, antagonize host cGAS DNA sensing [57]. Similar as their study, we found, siRNA containing a 5-nt motif antagonize host IFI16 DNA sensing, another important DNA sensor protein in host pathogen pattern recognition. In order to clarify whether the motif siRNA makes a complex with RIG-1 and IFI16, we performed motif siRNA-conjugated pull-down assay. The result as presented in the Figure EV2, indicated that motif siRNA (3’ or 5’) formed a complex with RIG-1 and IFI16, and IFI16 was not dissociated from the complex in the presence of DNA as a competitor, indicating that the persistence of the complex (RIG-1-motif siRNA-IFI16) hindered the binding of IFI16 to invaded DNA. A diagram of our findings is presented in the Figure 8.

**Figure 8.**
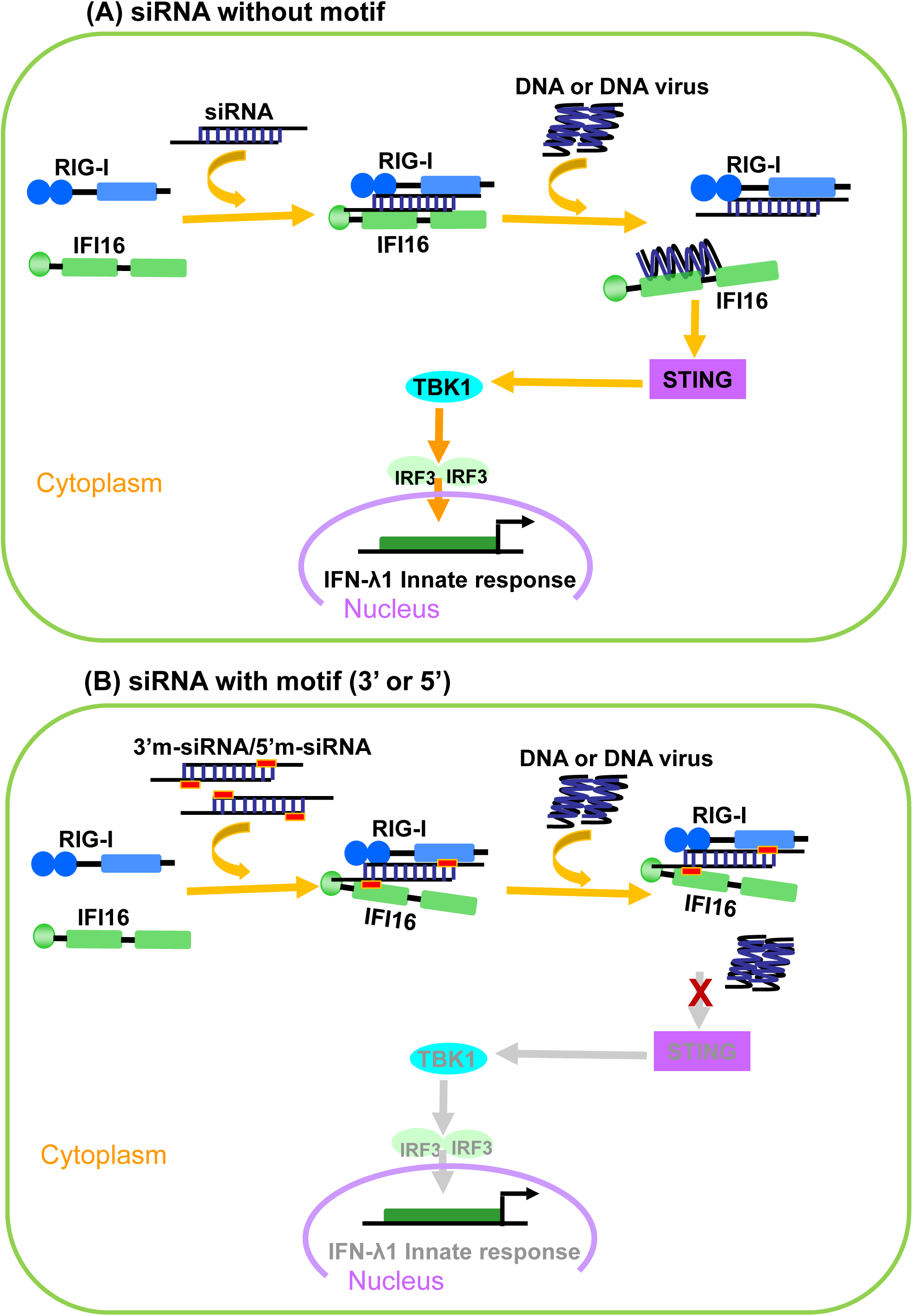
A simplified view of motif siRNA (3’ or 5’) inhibits IFI16-mediated innate immune response. (**A**) Ctrl-siRNA formed a complex with RIG-1 and IFI16. And IFI16 dissociated from the complex and bound with invaded DNA and therefore activated downstream STING-IRF3 signaling pathway. (**B**) Compared with Ctrl-siRNA, motif siRNA with the motif located at 3’ or 5’ terminus formed a similar RIG-1-siRNA-IFI16 complex. However, IFI16 didn’t dissociate from this complex to bind with invaded DNA, therefore blocking the activation of downstream signaling.

The sensing of self and foreign DNA is delicately regulated [59-61], and the dysregulation of DNA sensing is associated with several human autoimmune diseases, such as systemic lupus erythematosus and Aicardi-Goutières syndrome [62, 63]. Therefore, further delineation of the mechanisms by which to regulate the over-response of host cells may facilitate the development of therapeutics to treat or prevent diseases in which this pathway is dysregulated. Taken advantage of this broad anti-inflammatory and immunomodulatory properties, we reasoned that motif siRNA may prove to be beneficial as adjunctive therapy for autoimmune diseases.

## Materials and Methods

### Cell culture

Human cervical cancer (HeLa), SV40 T-antigen transformed human embryonic kidney cell 293 (293T) and acute human monocytic leukemia (THP-1) cell lines were obtained from American Type Culture Collection (ATCC) and maintained following manufacturer’s instructions. And THP1-Lucia ISG cells were purchased from InvivoGene (San Diego, CA) and maintained following the manufacturer’s protocol. To obtain human monocyte-derived immature dendritic cells (iDC), CD14+ monocytes were isolated from peripheral blood mononuclear cells (PBMCs) of healthy donors as previously described [21], and monocytes were cultured at 0.5 × 10^6^ cells/ml in RPMI1640 (Thermo Fisher Scientific, Waltham, MA) containing 10% fetal bovine serum (GE Healthcare, Pittsburgh, PA); 50 ng/ml Granulocyte-macrophage colony-stimulating factor (R&D Systems, Minneapolis, MN); 50 ng/ml IL-4 (R&D Systems); 25 mM HEPES (Quality Biology, Gaithersburg, MD) and 5 μg/ml Gentamicin (Thermo Fisher Scientific) for 7 days. Non-adherent cells were harvested as iDCs for use in the subsequent experiments.

### siRNAs

All siRNAs were chemically synthesized by Thermo Fisher Scientific or Integrated DNA Technologies (Coralville, IA).

### Plasmids

The pCR2.1 vector (Thermo Fisher Scientific) was digested by EcoR I (NEB, Ipswich, MA) and purified. This linearized plasmid was used to stimulate cells by transfection. The His-tagged IFI16 expression vector was constructed as previously described[21].

### Viruses and virus Infection

HSV-1 (MacIntyre strain) and Sendai virus (SeV) were obtained from Advanced Biotechnologies Inc. (Eldersburg, MD). HSV-1 virus was used at a MOI of 5, and SeV was used at a final concentration of 40 HA units/ml.

### Transfections

siRNAs were transfected into cells using Lipofectamine RNAiMAX (Thermo Fisher Scientific) according to the manufacturer’s protocols. DNA transfections were conducted with Lipofectamine 2000 (Thermo Fisher Scientific) for HeLa and THP-1 cells or TransIT-293 (Mirus Bio LLC, Madison, WI) for 293T cells, according to the manufacturers’ protocols. LyoVec (InvivoGen) was used for transfecting DNA agonists into THP-1 Lucia ISG cells.

### RNA extraction and real-time RT-PCR

Total cellular RNA was isolated from cells using the RNeasy isolation kit (Qiagen, Germantown, MD). The cDNA was synthesized from total RNA using Taqman reverse transcription reagents (Thermo Fisher Scientific) with random hexamer primer, according to the manufacturer’s instructions. IFN or inflammatory cytokine mRNA expression levels were measured using quantitative RT-PCR on a CFX96 real-time system (BioRad, Hercules, CA); the two-temperature cycle of 95 °C for 15 s and 60 °C for 1 min (repeated for 40 cycles) was used. Relative quantities of the transcript were calculated using the ΔΔCt method with GAPDH as a reference. Normalized samples were expressed relative to the average ΔCt value for controls to obtain relative fold change in expression levels. The probes specific for IFN or inflammatory cytokines were purchased from Applied Biosystems (Thermo Fisher Scientific).

### Western blot

Cell lysates were prepared using RIPA buffer (Boston BioProducts, Ashland, MA) in the presence of protease inhibitor cocktail (Sigma-Aldrich, St. Louis, MO) and Halt phosphatase inhibitor cocktail (Thermo Fisher Scientific). Protein concentrations of the cell lysates were quantified using a BCA protein assay (Thermo Fisher Scientific) to ensure equal amounts of total protein were loaded in each well of NuPAGE 4–12% Bis-Tris Gel (Thermo Fisher Scientific). Proteins were transferred onto a nitrocellulose membrane and blotted with the appropriate antibodies, followed by the HRP-conjugated secondary antibodies, and the detection was performed using ECL plus Western blotting detection reagents (GE Healthcare).

### Oligonucleotide pull-down assay

Cytoplasmic extracts were generated by disrupting cells using hypotonic buffer (Active Motif, Carlsbad, CA) and pelleting the cell debris by centrifugation at 14000 × g for 10 min at 4 °C. A total of 1 mg cytoplasmic protein was incubated with 100 μL Dynabeads M-280 streptavidin (Thermo Fisher Scientific) conjugated with 5′-biotinylated siRNA or 100 bp linearized DNA. Precipitated oligonucleotide-protein complexes were washed extensively in protein binding buffer (20 mM HEPES, 50 mM KCL, 10% glycerol, 5 mM MgCL2 and 1 mM DTT). Proteins were eluted from the beads by boiling in NuPAGE LDS sample buffer (Thermo Fisher Scientific) and were separated on NuPAGE 4– 12% Bis-Tris gel (Thermo Fisher Scientific) and detected by antibodies using western blot.

### AlphaScreen IFI16 and siRNA binding assay

The binding affinity of siRNA for His-tagged IFI16 was determined by an amplified luminescent proximity homogenous assay (AlphaScreen) as described [21]. Briefly, the whole cell lysate from His-tagged IFI16 overexpressed HEK 293T cells was incubated with different concentration of biotinylated siRNA for 1 h at 37°C in buffer (50 mM Tris/pH7.4, 100 mM NaCl, 0.01% Tween20, 0.1% BSA) and subsequently incubated for 30 min at 25°C with Nickel Chelate acceptor beads (Perkin Elmer, Waltham, MA) and Streptavidin conjugated donor beads (Perkin Elmer). The resulting chemiluminescence emitting in the range of 520–620 nm correlates with the number and proximity of associated beads which is inversely correlated with the dissociation constant of donor (biotin-siRNA) and acceptor (His-IFI16). The assay was performed in wells of half area 96-well plates (Perkin Elmer). Plates were analyzed for emitted fluorescence with a multilabel reader Envision (Perkin Elmer).

### Microarray

Total cellular RNA was isolated from cells using the RNeasy isolation kit (Qiagen, Germantown, MD). RNA was quantitated and qualified with Nanodrop 1000 (Thermo Fisher Scientific) and Agilent Bioanalyzer RNA Nano 6000 chip (Agilent, Santa Clara, CA). cRNA synthesis, labeling and hybridization to the Human GeneArray (Thermo Fisher Scientific) were performed according to the manufacturer’s protocol. Genechip data analysis was performed using Partek Genomics Suite (Partek, Inc., St Louis, MO). Briefly, Raw expression values for each array were normalized using a quantile normalization procedure. And the significant differences in expression were determined by one-way ANOVA. Functional enrichment analysis was performed using the DAVID (the database for annotation, visualization and integrated discovery) bioinformatics resources (https://david.ncifcrf.gov/home.jsp).

### Statistical analysis

Results were representative of at least three independent experiments. The values were expressed as mean and SD of individual samples. Statistical significance was determined by the Student’s t test. A P < 0.05 was considered a significant difference between the experimental groups.

## Acknowledgements

Authors thank H. Clifford Lane for supporting and discussing this project. This project has been funded in whole or in part with federal funds from the National Cancer Institute, National Institutes of Health, under Contract No. HHSN261200800001E. The content of this publication does not necessarily reflect the views or policies of the Department of Health and Human Services, nor does mention of trade names, commercial products, or organizations imply endorsement by the U.S. Government. This research was supported [in part] by the National Institute of Allergy and Infectious Disease.

## Author contributions

H.S. and T.I. designed and performed experiments; H.S. and T.I. discussed and interpreted the data; J.Y. and X.H. performed microarray and analyzed data; Q.C. provided iDCs. H.S. and T.I. wrote the manuscript.

## Expanded view Figure and Table legends

**Table EV1. The sequences (sense strand) of the immunostimulatory siRNAs which enhance DNA-mediated *IFNL1* signaling.**

**Table EV2. The details of genes in the set of 51 that belong to each class are listed.**

**Figure EV1. an RNA secondary structure analysis for motif siRNAs with motif sequence at different location.** RNAfold tool in ViennaRNA Web Services (http://rna.tbi.univie.ac.at/cgi-bin/RNAWebSuite/RNAfold.cgi) was used for RNA secondary structure prediction.

**Figure EV2. 3’m-siRNA bound with RIG-1 and IFI16, however IFI16 didn’t dissociate from the complex (RIG-1-3’m-siRNA-IFI16) when DNA was added as a competitor.** 3’m-siRNA-conjugated pull-down assay was performed in the absence or presence of DNA as a competitor. Bound proteins on the beads were separated by SDS-PAGE, and western blot was performed using anti-RIG-1 and anti-IFI16 antibodies.

## References

1. Medzhitov R, Janeway C (2000) Innate Immunity. N Engl J Med 343: 338–344

2. Sharma S, Fitzgerald KA (2011) Innate Immune Sensing of DNA. PLoS Pathog 7: e1001310

3. Christensen MH, Paludan SR (2016) Viral evasion of DNA-stimulated innate immune responses. Cellular & Molecular Immunology 14: 4

4. Zhang Z, Yuan B, Bao M, Lu N, Kim T, Liu Y (2011) The helicase DDX41 senses intracellular DNA mediated by the adaptor STING in dendritic cells. Nat Immunol 12: 959–965

5. Unterholzner L, Keating SE, Baran M, Horan KA, Jensen SB, Sharma S, Sirois CM, Jin T, Latz E, Xiao TS, et al. (2010) IFI16 is an innate immune sensor for intracellular DNA. Nat Immunol 11: 997–1004

6. Takaoka A (2007) DAI (DLM-1/ZBP1) is a cytosolic DNA sensor and an activator of innate immune response. Nature 448: 501–505

7. Yang P, An H, Liu X, Wen M, Zheng Y, Rui Y, Cao X (2010) The cytosolic nucleic acid sensor LRRFIP1 mediates the production of type I interferon via a [beta]-catenin-dependent pathway. Nat Immunol 11: 487–494

8. Kim T, Pazhoor S, Bao M, Zhang Z, Hanabuchi S, Facchinetti V, Bover L, Plumas J, Chaperot L, Qin J, et al. (2010) Aspartate-glutamate-alanine-histidine box motif (DEAH)/RNA helicase A helicases sense microbial DNA in human plasmacytoid dendritic cells. Proceedings of the National Academy of Sciences 107: 15181–15186

9. Zhang Z, Kim T, Bao M, Facchinetti V, Jung Sung Y, Ghaffari Amir A, Qin J, Cheng G, Liu Y-J (2011) DDX1, DDX21, and DHX36 Helicases Form a Complex with the Adaptor Molecule TRIF to Sense dsRNA in Dendritic Cells. Immunity 34: 866–878

10. Gao D, Wu J, Wu Y-T, Du F, Aroh C, Yan N, Sun L, Chen ZJ (2013) Cyclic GMP-AMP Synthase Is an Innate Immune Sensor of HIV and Other Retroviruses. Science 341: 903–906

11. Sun L, Wu J, Du F, Chen XG, Chen ZJ (2013) Cyclic GMP-AMP Synthase Is a Cytosolic DNA Sensor That Activates the Type I Interferon Pathway. Science 339: 786–791

12. Fernandes-Alnemri T, Yu JW, Datta P, Wu J, Alnemri ES (2009) AIM2 activates the inflammasome and cell death in response to cytoplasmic DNA. Nature 458: 509–513

13. Hornung V (2009) AIM2 recognizes cytosolic dsDNA and forms a caspase-1-activating inflammasome with ASC. Nature 458: 514–518

14. Burckstuummer T (2009) An orthogonal proteomic-genomic screen identifies AIM2 as a cytoplasmic DNA sensor for the inflammasome. Nature Immunol 10: 266–272

15. Kotenko SV (2011) IFN-λs. Curr Opin Immunol 23: 583–590

16. Donnelly RP, Kotenko SV (2010) Interferon-Lambda: A New Addition to an Old Family. J Interferon Cytokine Res 30: 555–564

17. Ank N, West H, Bartholdy C, Eriksson K, Thomsen AR, Paludan SR (2006) Lambda Interferon (IFN-λ), a Type III IFN, Is Induced by Viruses and IFNs and Displays Potent Antiviral Activity against Select Virus Infections In Vivo. J Virol 80: 4501–4509

18. Ank N, Paludan SR (2009) Type III IFNs: New layers of complexity in innate antiviral immunity. Biofactors 35: 82–87

19. Syedbasha M, Egli A (2017) Interferon Lambda: Modulating Immunity in Infectious Diseases. Frontiers in Immunology 8

20. Zhang X, Brann TW, Zhou M, Yang J, Oguariri RM, Lidie KB, Imamichi H, Huang DD, Lempicki RA, Baseler MW, et al. (2011) Cutting Edge: Ku70 Is a Novel Cytosolic DNA Sensor That Induces Type III Rather Than Type I IFN. The Journal of Immunology 186: 4541–4545

21. Sui H, Zhou M, Chen Q, Lane HC, Imamichi T (2014) siRNA enhances DNA-mediated interferon lambda-1 response through crosstalk between RIG-I and IFI16 signalling pathway. Nucleic Acids Res 42: 583–598

22. Sui H, Zhou M, Imamichi H, Jiao X, Sherman BT, Lane HC, Imamichi T (2017) STING is an essential mediator of the Ku70-mediated production of IFN-λ1 in response to exogenous DNA. Science Signaling 10

23. Travar M, Vucic M, Petkovic M (2014) Interferon Lambda-2 Levels in Sputum of Patients with Pulmonary Mycobacterium tuberculosis Infection. Scand J Immunol 80: 43–49

24. Yan B, Chen F, Xu L, Wang Y, Wang X (2017) Interleukin-28B dampens airway inflammation through up-regulation of natural killer cell-derived IFN-γ. Scientific Reports 7: 3556

25. Murakawa M, Asahina Y, Kawai-Kitahata F, Nakagawa M, Nitta S, Otani S, Nagata H, Kaneko S, Asano Y, Tsunoda T, et al. (2017) Hepatic IFNL4 expression is associated with non-response to interferon-based therapy through the regulation of basal interferon-stimulated gene expression in chronic hepatitis C patients. J Med Virol 89: 1241–1247

26. Lu Y-F, Goldstein DB, Urban TJ, Bradrick SS (2015) Interferon-λ4 is a cell-autonomous type III interferon associated with pre-treatment hepatitis C virus burden. Virology 476: 334–340

27. Sui H-Y, Zhao G-Y, Huang J-D, Jin D-Y, Yuen K-Y, Zheng B-J (2009) Small Interfering RNA Targeting M2 Gene Induces Effective and Long Term Inhibition of Influenza A Virus Replication. PLoS ONE 4: e5671

28. Akira S, Uematsu S, Takeuchi O (2006) Pathogen Recognition and Innate Immunity. Cell 124: 783–801

29. Mogensen TH (2009) Pathogen Recognition and Inflammatory Signaling in Innate Immune Defenses. Clin Microbiol Rev 22: 240–273

30. Turner MD, Nedjai B, Hurst T, Pennington DJ (2014) Cytokines and chemokines: At the crossroads of cell signalling and inflammatory disease. Biochimica et Biophysica Acta (BBA) - Molecular Cell Research 1843: 2563–2582

31. Sledz CA, Williams BRG (2005) RNA interference in biology and disease. Blood 106: 787–794

32. Sioud M (2010) Recent advances in small interfering RNA sensing by the immune system. New Biotechnology 27: 236–242

33. Sioud M (2007) RNA interference and innate immunity. Advanced Drug Delivery Reviews 59: 153–163

34. Whitehead KA, Dahlman JE, Langer RS, Anderson DG (2011) Silencing or Stimulation? siRNA Delivery and the Immune System. Annual Review of Chemical and Biomolecular Engineering 2: 77–96

35. Schlee M, Hornung V, Hartmann G (2006) siRNA and isRNA: two edges of one sword. Molecular Therapy 14: 463–470

36. Marques JT, Williams BRG (2005) Activation of the mammalian immune system by siRNAs. Nat Biotech 23: 1399–1405

37. Sioud M (2005) Induction of Inflammatory Cytokines and Interferon Responses by Double-stranded and Single-stranded siRNAs is Sequence-dependent and Requires Endosomal Localization. J Mol Biol 348: 1079–1090

38. Reynolds A, Anderson EM, Vermeulen A, Fedorov Y, Robinson K, Leake D, Karpilow J, Marshall WS, Khvorova A (2006) Induction of the interferon response by siRNA is cell type– and duplex length–dependent. RNA 12: 988–993

39. Olejniczak M, Polak K, Galka-Marciniak P, Krzyzosiak WJ (2011) Recent Advances in Understanding of the Immunological Off-Target Effects of siRNA. Curr Gene Ther 11: 532–543

40. Zhang Z, Weinschenk T, Guo K, Schluesener HJ (2006) siRNA binding proteins of microglial cells: PKR is an unanticipated ligand. J Cell Biochem 97: 1217–1229

41. Wang Q, Nagarkar DR, Bowman ER, Schneider D, Gosangi B, Lei J, Zhao Y, McHenry CL, Burgens RV, Miller DJ, et al. (2009) Role of Double-Stranded RNA Pattern Recognition Receptors in Rhinovirus-Induced Airway Epithelial Cell Responses. J Immunol 183: 6989–6997

42. Hornung V, Guenthner-Biller M, Bourquin C, Ablasser A, Schlee M, Uematsu S, Noronha A, Manoharan M, Akira S, de Fougerolles A, et al. (2005) Sequence-specific potent induction of IFN-alpha by short interfering RNA in plasmacytoid dendritic cells through TLR7. Nat Med 11: 263–70

43. Veeranki S, Duan X, Panchanathan R, Liu H, Choubey D (2011) IFI16 Protein Mediates the Anti-inflammatory Actions of the Type-I Interferons through Suppression of Activation of Caspase-1 by Inflammasomes. PLOS ONE 6: e27040

44. Arakawa R, Bagashev A, Song L, Maurer K, Sullivan KE (2010) Characterization of LRRFIP1. Biochem Cell Biol 88: 899–906

45. Juliane L, Stefan R, Nikolaus D, Julia E, Karolin M, Vincent VL, Hortense S, Dje NGP, Stefan H, Trinad C, et al. (2008) IFNβ responses induced by intracellular bacteria or cytosolic DNA in different human cells do not require ZBP1 (DLM-1/DAI). Cellular Microbiology 10: 2579–2588

46. Sioud M (2011) Promises and Challenges in Developing RNAi as a Research Tool and Therapy. In RNA, Nielsen H (ed) pp 173–187. Humana Press

47. Meng Z, Lu M (2017) RNA Interference-Induced Innate Immunity, Off-Target Effect, or Immune Adjuvant? Frontiers in Immunology 8

48. Samuel-Abraham S, Leonard JN (2010) Staying on message: design principles for controlling nonspecific responses to siRNA. FEBS Journal 277: 4828–4836

49. Kalali BN, Kollisch G, Mages J, Muller T, Bauer S, Wagner H, Ring J, Lang R, Mempel M, Ollert M (2008) Double-Stranded RNA Induces an Antiviral Defense Status in Epidermal Keratinocytes through TLR3-, PKR-, and MDA5/RIG-I-Mediated Differential Signaling. J Immunol 181: 2694–2704

50. Goodchild A, Nopper N, King A, Doan T, Tanudji M, Arndt G, Poidinger M, Rivory L, Passioura T (2009) Sequence determinants of innate immune activation by short interfering RNAs. BMC Immunology 10: 40

51. Judge A, MacLachlan I (2008) Overcoming the Innate Immune Response to Small Interfering RNA. Hum Gene Ther 19: 111–124

52. Judge A, Sood V, Shaw J, Fang D, McClintock K, MacLachlan I (2005) Sequence-dependent stimulation of the mammalian innate immune response by synthetic siRNA. Nat Biotechnol 23: 457–62

53. Zheng B, Sui H, Lin Y (2014) siRNA compositions and methods for potently inhibiting viral infection. In Google Patents

54. Blanco P, Palucka AK, Pascual V, Banchereau J (2008) Dendritic cells and cytokines in human inflammatory and autoimmune diseases. Cytokine Growth Factor Rev 19: 41–52

55. Roger T, Lugrin J, Le Roy D, Goy G, Mombelli M, Koessler T, Ding XC, Chanson A-L, Reymond MK, Miconnet I, et al. (2011) Histone deacetylase inhibitors impair innate immune responses to Toll-like receptor agonists and to infection. Blood 117: 1205–1217

56. Vitali P, Scadden ADJ (2010) Double-stranded RNAs containing multiple IU pairs are sufficient to suppress interferon induction and apoptosis. Nat Struct Mol Biol 17: 1043–1050

57. Wu J-j, Li W, Shao Y, Avey D, Fu B, Gillen J, Hand T, Ma S, Liu X, Miley W, et al. (2015) Inhibition of cGAS DNA Sensing by a Herpesvirus Virion Protein. Cell Host & Microbe 18: 333–344

58. Chanput W, Mes JJ, Wichers HJ (2014) THP-1 cell line: An in vitro cell model for immune modulation approach. International Immunopharmacology 23: 37–45

59. Liang Q, Seo Gil J, Choi Youn J, Kwak M-J, Ge J, Rodgers Mary A, Shi M, Leslie Benjamin J, Hopfner K-P, Ha T, et al. (2014) Crosstalk between the cGAS DNA Sensor and Beclin-1 Autophagy Protein Shapes Innate Antimicrobial Immune Responses. Cell Host & Microbe 15: 228–238

60. Konno H, Konno K, Barber Glen N (2013) Cyclic Dinucleotides Trigger ULK1 (ATG1) Phosphorylation of STING to Prevent Sustained Innate Immune Signaling. Cell 155: 688–698

61. Ablasser A, Hemmerling I, Schmid-Burgk JL, Behrendt R, Roers A, Hornung V (2014) TREX1 Deficiency Triggers Cell-Autonomous Immunity in a cGAS-Dependent Manner. The Journal of Immunology 192: 5993–5997

62. Ahn J, Barber GN (2014) Self-DNA, STING-dependent signaling and the origins of autoinflammatory disease. Curr Opin Immunol 31: 121–126

63. Barber GN (2014) STING-dependent cytosolic DNA sensing pathways. Trends in Immunology 35: 88–93

